# Explaining flexible continuous speech comprehension from individual motor rhythms

**DOI:** 10.1101/2022.04.01.486685

**Authors:** Christina Lubinus, Anne Keitel, Jonas Obleser, David Poeppel, Johanna M. Rimmele

## Abstract

When speech is too fast, the tracking of the acoustic signal along the auditory pathway deteriorates, leading to suboptimal speech segmentation and decoding of speech information. Thus, speech comprehension is limited by the temporal constraints of the auditory system. Here we ask whether individual differences in auditory-motor coupling strength in part shape these temporal constraints. In two behavioral experiments, we characterize individual differences in the comprehension of naturalistic speech as function of the individual synchronization between the auditory and motor systems and the preferred frequencies of the systems. Obviously, speech comprehension declined at higher speech rates. Importantly, however, both higher auditory-motor synchronization and higher spontaneous speech motor production rates were predictive of better speech-comprehension performance. Furthermore, performance increased with higher working memory capacity (Digit Span) and higher linguistic, model-based sentence predictability – particularly so at higher speech rates and for individuals with high auditory-motor synchronization. These findings support the notion of an individual preferred auditory– motor regime that allows for optimal speech processing. The data provide evidence for a model that assigns a central role to motor-system-dependent individual flexibility in continuous speech comprehension.

## 1. Introduction

Speech comprehension relies on temporal processing, as speech and other naturalistic signals have a complex temporal structure with information at different timescales(1). The temporal constraints of the auditory system limit our ability to understand speech at fast rates(2,3). Interestingly, the motor system can under certain conditions provide temporal predictions that aid auditory perception(4,5). Accordingly, current oscillatory models of speech comprehension propose that properties of the auditory but also the motor system affect the quality of auditory processing(6,7). In two behavioral experiments, we investigate how the auditory, the motor system, and their synchronization shape individual flexibility of comprehending fast continuous speech.

Auditory temporal constraints have been observed as preferred rates of auditory speech(8,9) processing (but also of tones(10,11), and amplitude modulated sounds(11–14)) and explained in the context of neurocognitive models of speech perception. According to such proposals, humans capitalize on temporal information by dynamically aligning ongoing brain activity in auditory cortex to the temporal patterns inherent to the acoustic speech signal(15–18). By hypothesis, endogenous theta brain rhythms in auditory cortex partition the continuous auditory stream into smaller chunks at roughly the syllabic scale by tracking quasi-rhythmic temporal fluctuations in the speech envelope. This chunking mechanism allows for the decoding of segmental phonology – and ultimately linguistic meaning(15,18–20). The decoding of the speech signal is accomplished seemingly effortlessly within an optimal range centered in the traditional theta band(18), whereas comprehension deteriorates strongly for speech presented beyond ~9 Hz(2,3). While much research has focused on the apparent *stability* of the average acoustic modulation rate at the syllabic scale(8,9), the *flexibility* in speech comprehension(9,21), that is, what constitutes individual differences in understanding fast speech rates, is poorly understood.

The motor system, and neural auditory-motor coupling in particular, is a plausible candidate to facilitate individual differences in auditory speech processing abilities. Two arguments supporting this notion are the motor systems’ modulatory effect on auditory perception(22–24) and its susceptibility to training(25–27). While there is evidence suggesting that the auditory and speech motor brain areas are intertwined during speech comprehension(28–32), the extent to which speech motor processing modulates auditory processing is debated(5,33,34). Specifically, endogenous brain rhythms in both auditory(20,35) and motor(35,36) cortex have been observed to track the acoustic speech signal, and are characterized by preferred frequencies(19,37,38). In contrast to neural measures of preferred frequencies(37–39), here we used a behavioral estimate termed “preferred” or “spontaneous” rate. Furthermore, neural coupling between auditory and motor brain areas during speech processing(35,36,40,41) has been hypothesized to provide temporal predictions about upcoming sensory events to the auditory cortex(4,41–43). The precision of these predictions may be proportional to the strength of auditory-motor cortex coupling.

Auditory-motor cortex coupling strength varies across the population, as shown by recent work(6,10,40,44,45). Assaneo et al.(40) developed a behavioral protocol (spontaneous speech synchronization test; SSS-test) which quantifies the strength of auditory-to-motor synchronization during speech production in individuals. The authors reported that auditory-motor synchronization is characterized by a bimodal distribution in the population, classifying individuals into high versus low synchronizers. (The rejection of unimodality has been previously shown with large sample sizes(40, see also:,46).) Importantly, in addition to superior behavioral synchronization, high synchronizers have stronger structural and functional connectivity between auditory and speech motor cortices (see 40,Figure 3A and B). Thus, the SSS-test provides not only a behavioral measure but also approximates individual differences in neuronal auditory-motor coupling strength. We propose that the individual variability in auditory-motor synchronization, previously observed to predict differences in word learning(40), syllable detection(6), and rate discrimination(10), as well as the individual variability in preferred auditory and motor rate, predicts differences in an individuals’ ability to comprehend continuous speech at fast syllabic rates.

The influence of individual auditory-motor coupling strength on behavioral performance has so far been established for behavioral paradigms using rather basic auditory and speech stimuli, e.g. tones or syllables(6,10,40). The current study assesses its importance in a more naturalistic context: during the comprehension of continuous speech. This adds several layers of complexity. First, as speech unfolds over time, processing of continuous, i.e. longer and more complex, speech naturally demands more working memory capacity for maintenance and access to linguistic and context information(47). Second, rich linguistic context is used to derive linguistic predictions about upcoming words and sentences(48–51). When linguistic predictability of a sentence is high(52), speech comprehension is improved, even in adverse listening situations(53,54). Thus, similar to auditory-motor synchronization, linguistic predictability offers a compensatory mechanism when comprehension is difficult.

In summary, we investigate the role of auditory-motor synchronization with the SSS-test and the role of preferred rhythms of the auditory and motor systems for the individual flexibility of the comprehension of continuous speech. First, based on an established literature(3,18,55–57), we expected a decline in comprehension performance at syllabic rates beyond the theta range. Second, as a faciliatory effect of auditory-motor coupling on auditory processing has been observed(6,10,40), we hypothesized that individual differences in comprehension performance could be predicted by individual auditory-motor synchronization, with superior speech comprehension for high synchronizers. Such a faciliatory effect might be strongest in demanding listening situations, such as at fast syllabic rates(5,10). Third, while the consequences of potential individual variation in the preferred rates of the motor and auditory systems are not clearly understood, based on previous findings(35) we expected a systematic relation of both preferred auditory and motor rates with individual speech comprehension performance. Finally, we hypothesized that linguistic predictability and working memory span should positively affect speech comprehension. Similar to auditory-motor synchronization, we expected linguistic predictability to interact with syllabic rate, such that both systems would become stronger predictors for speech comprehension as syllabic rate increases.

## 2. Methods

Two behavioral experiments and a control experiment were conducted: Experiment 1 was performed in the laboratory and investigated the influence of the spontaneous speech motor production rate on speech comprehension performance. In Experiment 2 we aimed to understand the complex interplay of multiple variables during speech comprehension beyond the spontaneous speech motor production rate. To this end, we additionally measured participants’ preferred auditory rate, auditory-motor synchronization, and working memory capacity. Experiment 2 and the control experiment were online studies. All studies were approved by the local ethics committees (Experiment 1: committee of the School of Social Sciences, University of Dundee, UK (No. UoD-SoSS-PSY-UG-2019-88), Experiment 2 and control experiment: procedures were approved by the Ethics Council of the Max Planck Society (2017_12)).

### 2.1. Participants

Participants were English native speakers with normal hearing and no neurological or psychological disorders (Exp 1: N = 34, Exp 2: N = 82, Control: N = 39). Participation was voluntary. For a detailed description of participants, stimuli, exclusion criteria, and tasks please refer to Supplementary Methods, Figures 1–2, and Tables 1-2.

**Figure 1.**
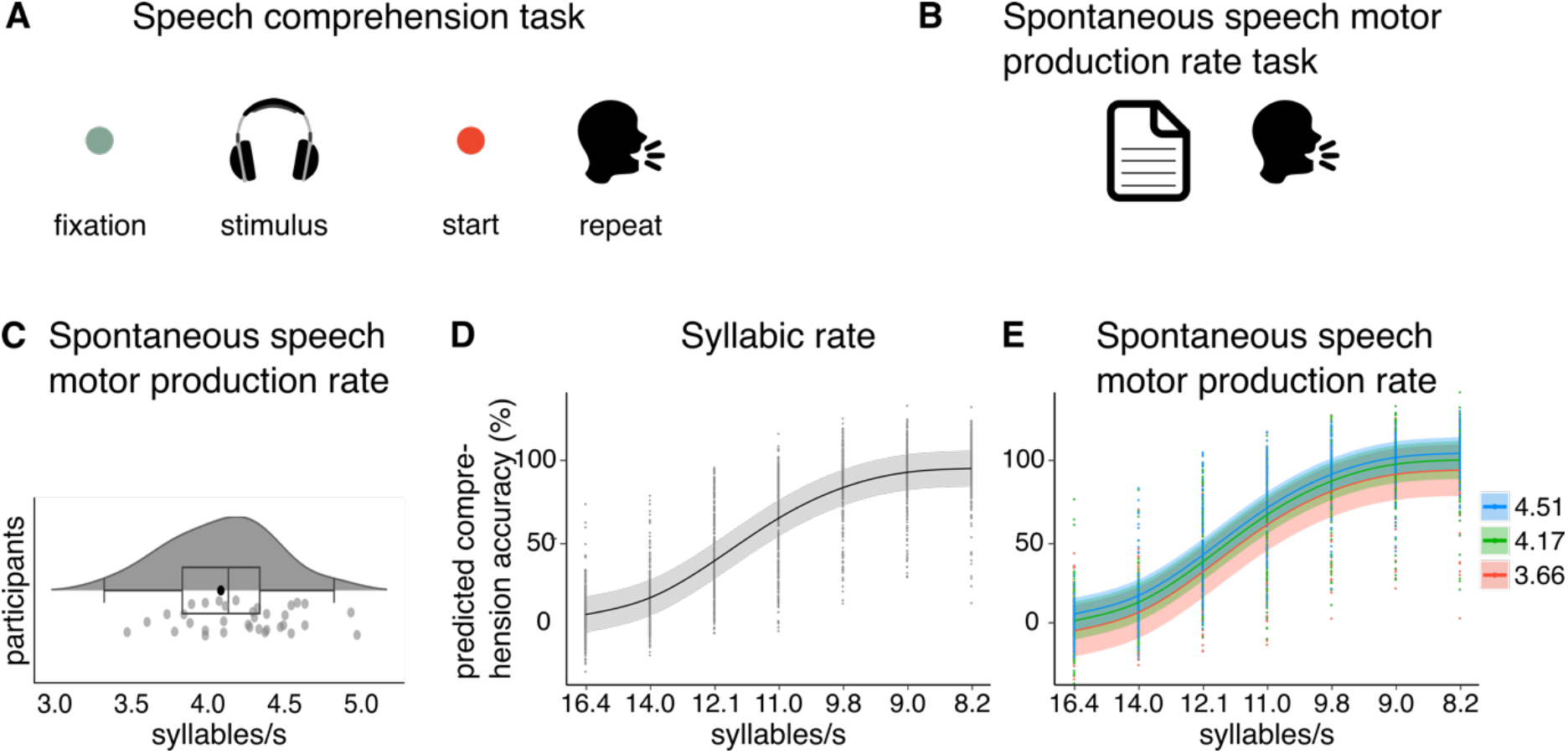
**A**. Example trial for the speech comprehension task. Participants fixated on a green fixation dot while presented auditorily with a sentence. On stimulus offset the fixation dot turned red, indicating to commence recall, i.e., reporting the sentence back. **B**. Spontaneous speech motor production rate task. Participants read a stimulus paragraph from a paper. **C**. Spontaneous speech motor production rate. We observed spontaneous speech motor production rates between 3.35 and 4.85 syllables/s (*M* = 4.11 syllables/s, left). The violin and boxplot show summary statistics and density: the median center line, 25^th^ to 75^th^ percentile hinges, whiskers indicate minimum and maximum within 1.5 × interquartile range. Grey dots represent participants individual speech motor productions rates, averaged across 6 trials. **D**. Main effect of syllabic rate. Plot shows the predicted main effect of syllabic rate from the generalized additive mixed model (GAMM). Black line indicates the predicted effect with 95% confidence interval in grey. Black dots show trial-level speech comprehension performance per subject and rate condition. **E**. Main effect of spontaneous speech motor production rate. Plot shows the predicted main effect of spontaneous speech motor production rate from the GAMM. Colored lines indicate the predicted effect with 95% confidence interval in the corresponding color. Colored dots show trial-level speech comprehension performance per subject and rate condition.

**Figure 2.**
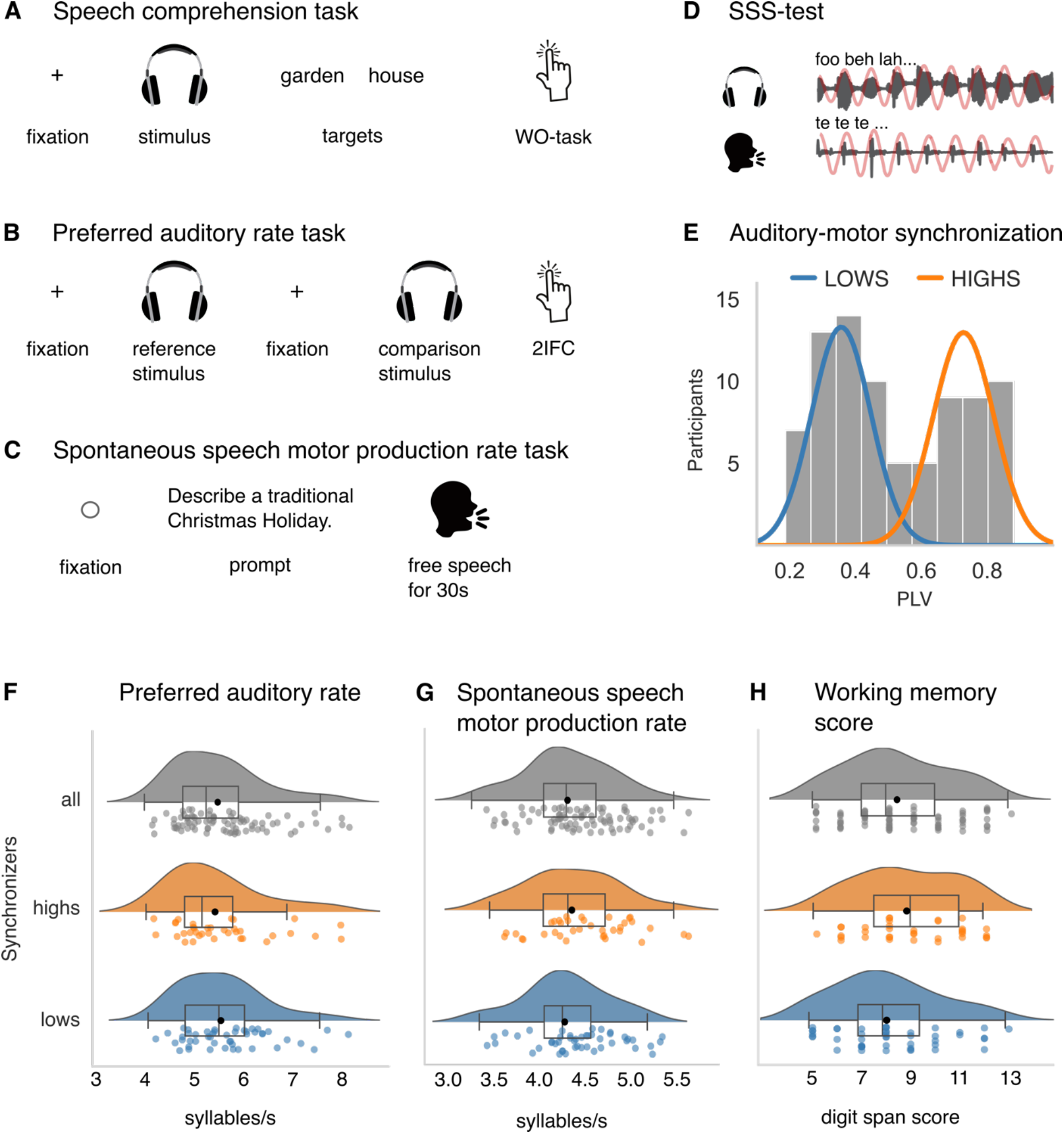
Panels **A, B**, and **C** visualize example trials for the speech comprehension task, the preferred auditory rate task, and the spontaneous speech motor production task, respectively. **D**. Schematic representation of the SSS-test, used to measure auditory-motor synchronization. Participants whisper a syllable (here /te/). **E**. Histogram of auditory-motor synchronization strength, obtained with the SSS-test. Participants were classified into high and low synchronizers (highs, lows) based on their PLV using k-means clustering. Group affiliation is overlaid by colored lines representing fitted normal distributions. **F**. Participants showed a mean preferred auditory rate of 5.57 syllables/s (*SD* = 0.86), with no differences between high and low synchronizers (*U* = 897.5, *p* = .48). **G**. We observed a spontaneous speech motor production rate between 3.36 and 5.38 syllables/s (*M* = 4.32, *SD* = 0.45) and no group difference between high and low synchronizers (*U* = 767.0, *p* = .60). **H**. Working memory capacity was indicated by a mean digit-span forward score of 8.46 (*SD* = 2.12) and the score did not differ between high and low synchronizers (*U* = 666.5, *p* = .14).

### 2.2. Design and materials

#### Speech comprehension task

In two speech comprehension tasks, we measured participants ability to comprehend sentences at various syllabic rates. Sentences were presented at 7 (Exp 1: [8.2, 9.0, 9.8, 11.0, 12.1, 14.0, 16.4]) or 6 (Exp 2: [5.00, 10.69, 12.48, 13.58, 14.38, 15.00]) rates. In Experiment 1, participants performed a classic intelligibility task, i.e. also termed “word identification task” (58,59, for review:,60). On each trial (N = 70), a sentence was presented through headphones and participants verbally repeated the sentence as accurately as possible (Fig. 1A). Responses were recorded.

In Experiment 2, speech comprehension was measured by a word-order task. Participants listened to one sentence per trial (N = 240), followed by the presentation of two words from the sentence on screen. Participants indicated via button press which word they heard first (Fig. 2A).

#### Speech production task

In the speech production tasks we estimated participants individual spontaneous speech motor production rate. In Experiment 1, the speech production task was operationalized by participants reading a text excerpt (216 words) from a printout. Participants were instructed to read the text excerpt out loud at a comfortable and natural pace while their speech was recorded (Fig. 1B).

In Experiment 2, participants were asked to produce continuous, “natural” speech. To facilitate fluent production, they were prompted by a question/statement belonging to six thematic categories (6 trials; own life, preferences, people, culture/traditions, society/politics, general knowledge, see Supplementary Table 2). Each response period lasted 30 seconds and trials were separated by self-paced breaks (Fig. 2C). While speaking, participants simultaneously listened to white noise. The white noise was introduced to measure the preferred rate of the motor system, without potential interference from auditory feedback. A second reason was to be consistent with the protocol from the SSS-test(40,61; also see below). Note that this procedure was not applied in Experiment 1.

#### Auditory rate task (only Exp 2)

To measure participants preferred auditory rate, we implemented a two-interval forced choice (2IFC) task, presenting a reference and a comparison stimulus at each trial. Participants indicated via button press which stimulus they preferred (Fig. 2B). Stimuli were presented at syllabic rates from 3.00 to 8.50 syllables/s (3.00, 3.92, 4.83, 5.75, 6.67, 7.58, 8.50). A reference rate, e.g. 3.00 syllables/s, was compared to all syllabic rates, including itself. For each reference/comparison pair the same sentence was presented – the stimuli only differed in their syllabic rates. Additionally, the task included catch trials to measure participant’s engagement (see Supplementary Methods for details).

#### Spontaneous speech synchronization (SSS) test (only Exp 2)

We measured participant’s auditory-motor synchronization using the SSS-test (for details:,40). In the main task, participants listened to a random syllable train and whispered along for a duration of 80s. They were instructed to synchronize their own syllable production to the stimulus presented through their headphones (Fig. 2D). The syllable rate in the auditory stimulus progressively increased in frequency from 4.3 to 4.7 syllables/s in increments of 0.1 syllables/s, every 60 syllables. Participants completed two trials, while the whispering was recorded.

Participants’ syllable production was masked by the simultaneously presented auditory syllable train. The masking procedure suppresses auditory feedback, allowing us better to isolate the synchronization of motor production to the auditory input, without interference of auditory feedback(44).

#### Digit span test (only Exp 2)

Working memory capacity was quantified using the forward and backward(62) digit span test. As for the backward test data is missing for N = 21 participants, only the forward span is reported. Digit spans were presented auditorily and participants typed in their responses(63).

#### Control Experiment

We designed a control experiment to test if the correct word order from the word order task of Experiment 2 could be guessed from the target words alone, that is, without understanding the sentence. The task consisted in judging which of two words would be more likely to occur first in a *hypothetical sentence*. On each trial, two words were presented on screen and participants indicated their choice via button press. Importantly, 1) participants did not listen to a full sentence at any time and 2) the target words were taken from the stimulus materials actually presented in Experiment 2.

### 2.3. Analysis

#### Spontaneous speech motor production rate (Exp 1 + 2)

The individual *spontaneous speech motor production rate*, i.e. articulation rate(64), was computed using Praat software(65) by automatically detecting syllable nuclei. The number of syllable nuclei was divided by the duration of the utterance, disregarding silent pauses. For Experiment 1, the production rate was computed across the entire reading paragraph. For Experiment 2, it was first calculated for each trial (30 s) separately. The motor rate was then averaged across all trials.

#### Preferred auditory rate (Exp 2)

First, participants with low performance in the catch trials of the preferred auditory rate task (below 75% correct) were excluded; amongst the remaining participants (*N* = 82) catch trial performance was very high (*M* = 98.48%, *SD* = 3.71). To compute the preferred auditory rate, a distribution of preferred frequencies was derived from all trials -except catch trials-by aggregating the frequency of each trials’ preferred item. Then a gaussian function was fitted to each participants’ distribution and two parameters were extracted: the peak as index for the preferred frequency and the full-width-at-half-maximum (FWHM) as index for the specificity of the response (lower FWHM equals stronger preference for one frequency).

#### Auditory-motor synchronization (Exp 2)

From the SSS-test(40) we derived the participant’s auditory-motor synchronization by calculating the phase-locking value (PLV)(66) between the (cochlea) envelopes of the auditory and the speech signals.

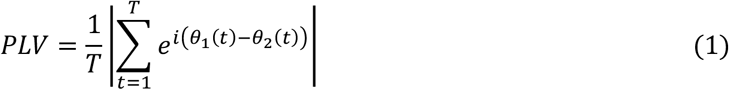

where *T* is the total number of time points, *t* denotes the discretized time, and *θ_1_* and *θ_2_* are the phase of the first and the second signals, respectively.

To obtain the cochlear envelope of the syllable train (auditory channels: 180–7,246 Hz), we used the Chimera Software toolbox(67). For the recorded speech signal the amplitude envelope was quantified as the absolute value of the Hilbert transform. Both envelopes were downsampled to 100 Hz and bandpass filtered (3.5-5.5 Hz) before their phase was extracted by means of the Hilbert transform. The PLV was first estimated for each trial of the SSS-test (time windows 5s, overlap 2s) and then averaged across runs, resulting in a mean PLV. The distribution of mean PLV values was subjected to a k-means algorithm(68) (*k* = 2) to split participants into a high- and a low-synchronizer group. Speech auditory-motor synchronization (PLV) was treated as bimodal variable based on previous research that rejected unimodality based on larger samples(40, see also:,46).

#### Linguistic predictability – Recurrent neural network (Exp 2)

Linguistic predictability of all stimulus sentences was measured by deriving single-sentence perplexity from a recurrent neural network language model. A language model, such as a recurrent neural network, assigns probabilities to all words in a sequence of words. From the single-word probabilities, we derived one value per sentence, quantifying its predictability(69,70). This so-called perplexity is the most common intrinsic evaluation metric of language models(71–73). It is computed as the inverse of the mean probability of a sentence weighted by sentence length(69), i.e. lower perplexity values equal higher sentence predictability (see Supplementary Methods for full details on RNN and perplexity).

#### Mixed effects models

For both experiments we performed mixed effects analyses to quantify how speech comprehension was affected by all variables of interest. Mixed models were computed using the R packages *lme4* (v1.1-29) and *mgcv* (v1.8-39), as set up in Rstudio (version 2022.2.1.461). Mixed-effects, rather than fixed-effects models were chosen to account for idiosyncratic variation within variables, i.e. repeated measures and therefrom resulting interdependencies between data points(74,75). Thus, both models included random intercepts for *participant* and *items*.

In Experiment 1, we computed a generalized additive mixed-effects model (GAMM) using the *mgcv:gam* function. For the dependent variable *speech comprehension*, we calculated the percentage of correctly repeated words for each sentence and subject from the speech comprehension task. The number of correct words was counted manually and transformed into a percentage. Then the dependent variable (single-trial data) was modelled as a function of the fixed effects *syllabic rate* and *spontaneous speech motor production rate*. A random slope for *syllabic rate* could not be included because the model failed to converge, thus the model included only random intercepts. Overall, the model explained ~77% of the variance.

In Experiment 2, the dependent variable *speech comprehension* was binary (*correct* vs *incorrect* word order judgment). Thus, we employed a generalized linear mixed-effects model (GLMM; *lme4:glmer* function) with a binomial logit link function. In terms of fixed effects, the model included all variables of interest: *syllabic rate*, *preferred motor rate*, *preferred auditory rate*, *auditory-motor synchronization*, *working memory, sentence predictability*. Additionally, we introduced several linguistic and other covariates for nuisance control(76): *predictability target 1*, *predictability target 2*, *sentence length* (# of words), *target distance* (i.e., distance in words between the target words), *compression/dilation* of audio file. In addition to random intercepts, the model contained a by-participant random slope for *syllabic rate*, allowing the strength of the effect of the rate manipulation on the dependent variable to vary between participants(74,75). Continuous predictor variables were z-transformed to facilitate the interpretation and comparison of the strength of the different predictors(77). Thus, the coefficients of all continuous predictors reflect log changes in comprehension for each unit (*SD*) increase in a given predictor. We observed no problems with (multi-)collinearity, all variance inflation factors were < 1.2 (package car version 3.0-10(78)). Overall, the model explained ~38% of the variance.

#### Control experiment

For each trial, we computed how many participants correctly guessed the word order (in percent, “*word order index*”). In a new GLMM analysis, this *word order index* was added as co-variate into the model from the main analysis while all other parameters remained the same.

## 3. Results

### 3.1. Experiment 1

In Experiment 1, we asked the question: to what extent is speech comprehension affected by one’s spontaneous speech motor production rate? *Speech comprehension* was measured as the percentage of correctly repeated words in an intelligibility task (2.75% to 93.70% on average across participants). We observed a mean *spontaneous speech motor production rate* of 4.11 syllables per second (*SD* = 0.35, min = 3.35, max = 4.85) across participants (Fig. 1C).

As expected, the GAMM revealed a main effect of *syllabic rate*: slower speech stimuli were associated with better speech comprehension (edf = 4.61, *F* = 1260.90, *p* < .001, Fig. 1D, see Supplementary Table 3). Importantly, we observed that the *spontaneous speech motor production rate* influenced speech comprehension: the higher the individual *spontaneous speech motor production rate*, the better the speech comprehension performance (edf = 1.00, *F* = 4.37, *p* = .036, Fig. 1E).

### 3.2. Experiment 2

First, in line with the first experiment, we observed a mean *spontaneous speech motor production rate* of 4.32 syllables per second across participants (*SD* = 0.45, Min = 3.36, Max = 5.38 syllables per second, Fig. 2G). Within-subject variance was low (Supplementary Fig. 3), suggesting that participants’ articulation rate was stable across trials. Second, participants showed a *preferred auditory rate* of ~5.57 syllables per second (peak: *M* = 5.57, *SD* = 0.86, Min = 4.16, Max = 7.92; FWHM, *M* = 4.89, *SD* = 0.50, Min = 3.23, Max = 5.50; Fig. 2F). Single-subject raw data can be inspected in Supplementary Fig. 4. Third, *auditory-to-motor speech synchronization* was quantified using the SSS-test(40), classifying participants as HIGH or LOW synchronizers (mean PLV HIGHs = 0.73, *SD* = 0.09, mean PLV LOWs = 0.36, *SD* = 0.09, Fig. 2E). Fourth, *working memory* was measured by means of the digit span test (62) which revealed a mean forward digit score of *M* = 8.46 (*SD* = 2.12, Min = 5.00, Max = 13.00, Fig. 2H).

The GLMM revealed that *syllabic rate* significantly influenced participants’ comprehension accuracy: for each increase of syllabic rate by one syllable/s, the odds of a correct word order judgment decreased (*odds ratio (OR)* = 0.65, *std. error (SE)* = 0.04, *p* < .001, Fig. 3A). This main effect of syllabic rate is consistent with a decline of speech comprehension performance at higher syllabic rates (3). In line with our hypothesis, we observed main effects for *spontaneous speech motor production rate* and *auditory-motor synchronization*. The higher a participant’s *spontaneous speech motor production rate*, the better the performance in the word order task (*OR* = 1.19, *SE* = 0.09, *p* = .014, Fig. 3C), replicating our finding from the first experiment. For *auditory-motor synchronization*, being a dichotomous variable (i.e., HIGH vs. LOW)(40), performance in the word order judgment task was higher for high compared to low synchronizers (*OR* = 1.34, *SE* = 0.20, *p* = .048, Fig. 3B). That is, across all trials, high synchronizers were more likely to correctly perform the task. Additionally, the model revealed a positive effect for *working memory score* (*OR* = 1.20, *SE* = 0.09, *p* = .012, Fig. 3D). This main effect suggests that better working memory performance enabled participants to better perform on the speech comprehension task. We did not observe a reliable effect of *preferred auditory rate* on speech comprehension (*OR* =1.14, *SE* = 0.08, *p* = .072). In contrast to our hypothesis, we observed no interaction effect of *syllabic rate* and *auditory-motor synchronization* on speech comprehension (*OR* = 0.97, *SE* = 0.07, *p* = .602).

**Figure 3.**
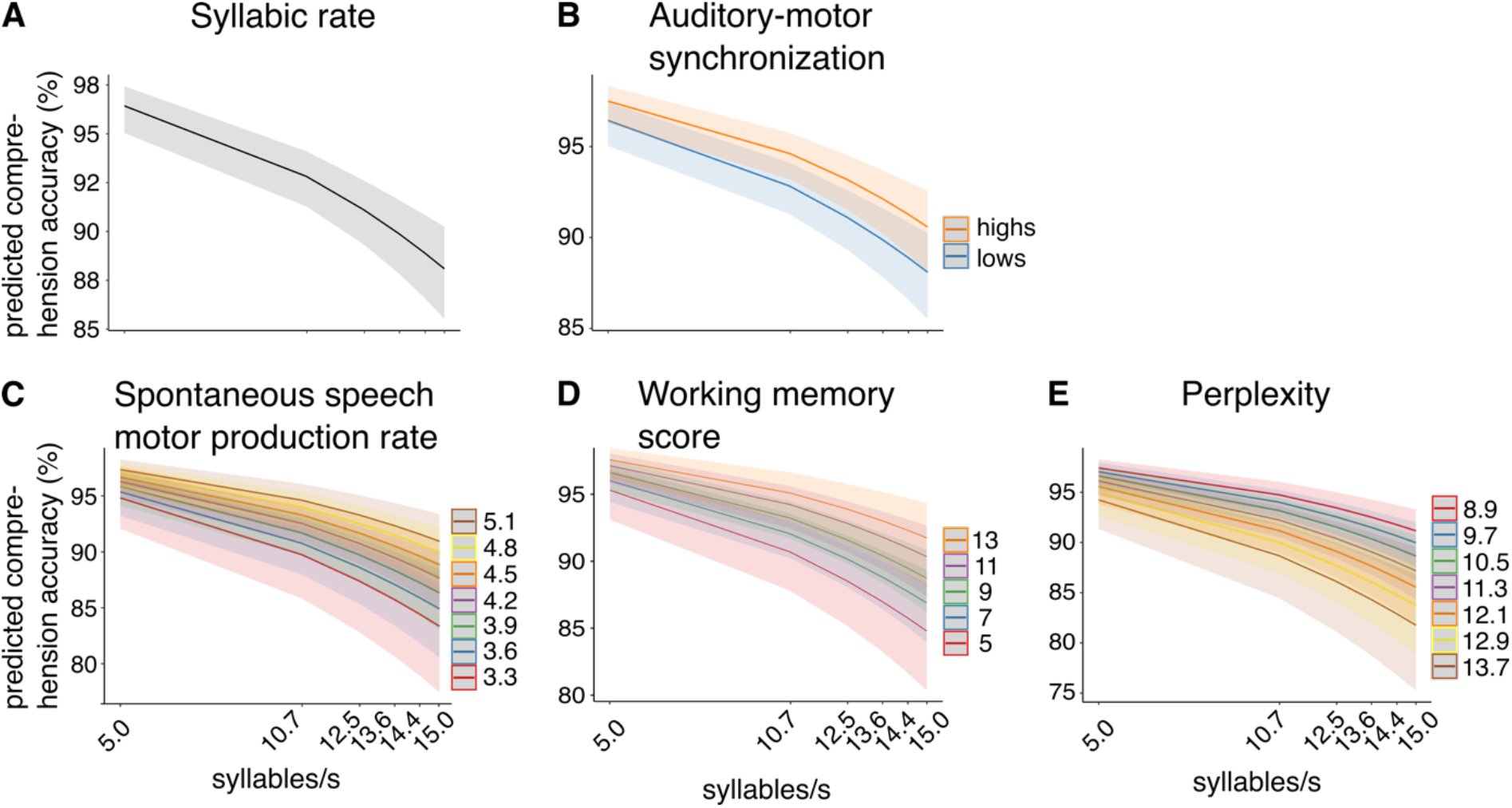
Significant main effects predicting speech comprehension performance. The generalized linear mixed effects model revealed a negative main effect of syllabic rate (A) and positive main effects of auditory-motor synchronization (B), spontaneous speech motor production rate (C), and working memory score (D). For stimulus perplexity we observed a negative main effect (E). In all panels, error shades indicate 95% confidence intervals. Note that the predictors are shown as a function of syllabic rate for visualization purposes only.

#### Linguistic predictability and further linguistic variables

To account for the effect of linguistic attributes, we expanded the GLMM by adding several (information-theoretic) linguistic variables: *perplexity*, *probability of target words*, *target distance*, and *stimulus length*. Adding these variables (with linguistic variables, AIC: 12675) improved model fit (without linguistic variables, AIC: 12848), as measured by a likelihood ratio test (χ2(6) = 184.24, p < .001, see Supplementary Table 4).

The full GLMM revealed that *perplexity* had a statistically reliable, negative effect on speech comprehension (*OR* = 0.84, *SE* = 0.04, *p* = .001, Fig. 3E) such that sentences with lower perplexity (which is equal to higher sentence predictability) lead to better speech comprehension performance. Additionally, we observed significant negative effects for *probability of target word 1* (*OR* = 0.93, *SE* = 0.03, *p* = .026) and *target word 2* (*OR* = 0.92, *SE*: 0.03, *p* = .021). Contrary to the perplexity effect, this suggests that task performance in the comprehension task was increased for unexpected target words.

Furthermore, the model revealed a positive effect for *target distance* (*OR* = 1.48, *SE*: 0.05, *p* < .001), suggesting that larger distance between targets was associated with better speech comprehension performance. In contrast, suggesting the opposite relation, for *stimulus length* we observed a negative effect (*OR* = 0.61, *SE*: 0.03, *p* < .001), i.e., shorter sentences resulted in higher comprehension performance. Due to the large number of variables introduced for nuisance control, we applied a control for multiple comparisons (i.e. false discovery rate; for full results see Supplementary Table 5). All effects remained robust after FDR correction: syllabic rate: *p* < .001; spontaneous speech motor production rate: *p* = .023; preferred auditory rate: *p* = .078; working memory score: *p* = .022; perplexity: *p* = .003; probability target 1: *p* = .034; probability target 2: *p* = .030; compression: *p* < .001; sentence length: *p* < .001; target distance: *p* < .001. Only auditory-motor synchronization changed from a significant effect to a trend (*p* = 0.057) (Note that this was a planned comparison and therefore is discussed).

Finally, we explored interaction effects between *syllabic rate*, *auditory-motor synchronization*, and *perplexity*. Adding the interaction term improved model fit (*χ^2^* (3) = 13.84, *p* = .004 (AIC without interaction term: 12675, AIC with interaction term: 12668)). The model revealed two significant 2-way interaction effects: *syllabic rate* × *perplexity* (*OR* = 0.88, *SE* = 0.05, *p* = .015) and *auditory-motor synchronization* × *perplexity* (*OR* = 0.86, *SE* = 0.04, *p* = .003; see Supplementary Fig. 5 and Supplementary Table 6). The interaction effect between *syllabic rate* and *perplexity* indicates that particularly comprehension of sentences at fast syllabic rates improves when *perplexity* is low. Furthermore, the *auditory-motor synchronization* × *perplexity* interaction effect suggests that while having better overall speech comprehension, high synchronizers show a stronger effect of *perplexity* compared to low synchronizers, with even better speech comprehension for more predictable sentences. The *syllabic rate* × *auditory-motor synchronization* effect (*OR* = 0.94, *SE* = 0.07, *p* = .392), as tested before, and the three-way interaction effect of *syllabic rate* × *auditory-motor interaction* × *perplexity* (*OR* = 1.09, *SE* = 0.06, *p* = .106) did not show a statistically reliable effect on speech comprehension.

#### Control experiment

In Experiment 2, speech comprehension performance was exceptionally good, even at high syllabic rates. To ensure the high performance was not an artifact of the task or stimuli, we conducted a control experiment. The analysis revealed that *word order index* did not influence speech comprehension in a statistically meaningful way (*OR* = 0.96, *SE* = 0.07, *p* = .219, see Supplementary Table 7).

## 4. Discussion

In two behavioral experiments, we show clear effects of *syllabic rate* on the comprehension of continuous speech. This finding is in line with proposals of speech comprehension being temporally constrained such that it is optimal for speech at lower syllabic rates. Crucially, in both protocols we observed that speech comprehension across a wide range of frequencies (5-15 syllables/s) was affected by participants’ *spontaneous speech motor production rate*, with higher rates predicting better speech comprehension. In the second experiment we showed that, beyond the spontaneous rate of the speech motor system, the individual strength of speech auditory-motor synchronization also affected comprehension. In contrast, the preferred speech perception rate was not related to speech comprehension performance. Together, these findings suggest that while speech comprehension is limited by general processing characteristics of the auditory system, interindividual differences in comprehension flexibility are moderated by the motor system and interactions between the auditory and motor systems (Fig. 4). Our findings furthermore allow us to generalize the effects of individual differences in the motor system on auditory perception, which have been previously shown for simpler stimuli(6,10,40,79), to more natural continuous speech.

**Figure 4.**
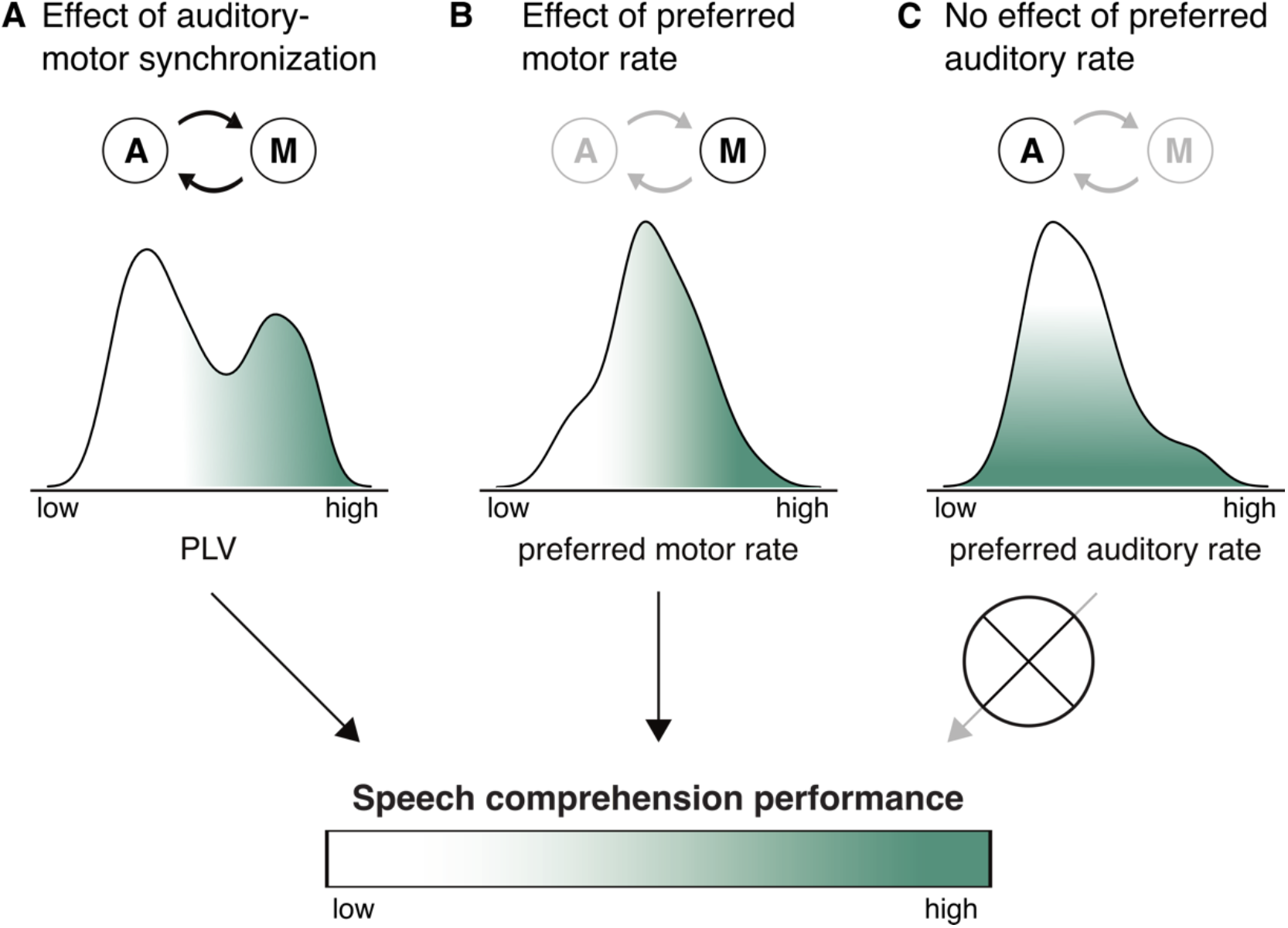
Schematic illustrating the relationship between speech comprehension performance and auditory-motor synchronization (A), the preferred rates of the motor (B) and of the auditory systems (C). All three predictor variables are represented by the corresponding distribution generated from our experimental data. We propose that better speech comprehension at demanding rates - and by hypothesis auditory behavior more generally - is accompanied by a higher preferred rate of the motor system as well as stronger auditory-motor synchronization. In contrast, the preferred rate of the auditory system seems not to determine auditory behavior. Circled A and M illustrate the auditory and motor systems. The arrows connecting them express the relevance of synchronization between the systems for the variable in question.

As expected(2,18,55–57), we observed that speech comprehension accuracy declined as syllabic rate increased. Although speech comprehension dropped at higher rates in both paradigms, the overall level of comprehension accuracy was much higher in Experiment 2, with accuracy remaining very high (~85%), even for speech as fast as 15 syllables/s. In contrast, in Experiment 1 the increase in syllabic rate resulted in a dramatic drop of comprehension performance. This is in line with our expectations, as the nature of the word-order task is likely to yield overall better performance than the classic intelligibility task. Additionally, our control experiment rules out a potential confound by demonstrating that the high performance in Experiment 2 is not due to simple guessing of the correct word order (see Results section and Supplementary Table 7). Interestingly, however, in both experiments performance decreased later than previously observed, that is, beyond rates of 9 syllables/s(56,80). However, in line with our findings, several other studies, also observed shallower decreases in speech comprehension, with relatively high comprehension at higher syllable rates (~12 syllables/s)(3,56,81,82). We consider several possible explanations for these discrepancies. One explanation for the different and higher speech-rate decline in comprehension performance is that naturally produced fast speech (with matched degrees of compression across syllabic rates, as used in Experiment 2), in contrast to linearly compressed speech, results in more variance of the speech rate and thus allows for part of the sentences to be understood. However, this explanation does not account for Experiment 1, in which all stimuli were synthesized at the same rate (varying in degrees of compression). Furthermore, the high performance level might be related to different complexity between more naturalistic sentences, providing stronger context information to compensate loss of information, as compared to the words(18), digits(83), or simple sentences(55) used in previous work. Finally, it is notable that while some studies conceptualized the syllabic rate based on the ‘theta-syllable’ (an information unit defined by cortical function(84)), we define syllabic rate as linguistically defined syllables per second, following other studies(36).

Auditory-motor speech synchronization, a behavioral estimate of auditory-motor cortex coupling strength(40), had a modulatory -albeit small-effect on speech comprehension. We observed that high compared to low synchronizers exhibited better speech comprehension performance. These results expand on findings which showed superior statistical word learning(40) or syllable discrimination(6) for individuals with stronger auditory-motor coupling by showing a similar effect for comprehending more naturalistic, continuous speech. Note that this effect requires further validation as it did not survive control for multiple comparisons (Supplementary Table 5). Additionally, we expected an interaction of syllabic rate and auditory-motor synchronization, as reported for rate discrimination in tone sequences(10). However, the modulation observed here occurred across all syllabic rates, suggesting that an interaction effect may be masked and compensated for by context and linguistic information in continuous speech comprehension. Alternatively, it is possible -although unlikely-that the interaction of syllabic rate and auditory-motor synchronization was not observed here due to the different frequency resolution at low frequencies. The difference between HIGHs and LOWS in Kern et al.(10) manifested between 7.14 and 10.29 Hz. In contrast, in the present experiment, there was no frequency condition between 5 and 10.69 syllables/s.

Importantly, the spontaneous motor production rate affected speech comprehension, suggesting that individuals with higher spontaneous motor production rate have increased speech comprehension abilities (at the higher range). We replicated this finding in the second experiment. The finding might reflect a complex interplay of auditory and motor cortex during speech comprehension wherein not only the coupling strength, but also the preferred rates of the motor cortex affect speech perception. A possible role of the preferred speech motor rate for speech processing has been previously discussed(35). Furthermore, our findings are in line with an oscillatory model of speech comprehension(6). An alternative interpretation of our findings might be that general processes such as vigilance and fatigue are equally reflected in the spontaneous speech motor production rate and the speech comprehension performance. This could be, because speech comprehension naturally is tightly intertwined with production, and vigilance effects on production for example might similarly affect comprehension. The behavioral protocol does not allow to completely discard such an alternative interpretation. However, given that no correlation of a demanding cognitive task (Digit span) with the spontaneous speech motor production rates was observed (see Supplementary Material), we consider this unlikely. Furthermore, for the effects of speech auditory-motor synchronization on syllable discrimination others have ruled out such an interpretation(6).

Interestingly, the preferred auditory rate (~5.55 syllables/s) had no effect on speech comprehension in our study. A possible explanation is that preferred rates in auditory cortex are less flexible compared to preferred rates in motor cortex and thus less prone to individual difference related improvements of speech comprehension. However, comparing the variances of the distribution of preferred auditory (*s^2^* = 0.74) and motor (*s^2^* = 0.20) rates revealed bigger variance in the auditory rate (*F*(1, 162) = 22.39, *p* < .001). Another possibility is that the behavioral estimation of preferred auditory cortex rates were not optimally operationalized. This might also explain the lack of correlation between preferred auditory and spontaneous speech production rates (see Supplementary Material), which we expected to be correlated. Generally, our behavioral protocol only allows for an indirect assessment of preferred neural rates. Nevertheless, behavioral measures have been regarded as proxy for underlying intrinsic brain rhythms(45). Finally, the rates at which speech comprehension decreases are much higher than the preferred auditory and spontaneous speech motor production rates. While the preferred rates were well within the expected range(7,8), the mismatch between maximal comprehension rates and preferred rates was due to the high speech comprehension ability of participants even at high rates.

We show that continuous speech comprehension is additionally affected by other higher cognitive and linguistic factors. The relevance of linguistic predictability and working memory capacity have been shown in multiple studies(53,54). In agreement with these studies, such cognitive variables explained a large amount of variance in speech comprehension. Interestingly, our findings suggest that the faciliatory effect of linguistic predictability is particularly effective at fast rates. Second, we tentatively interpret that facilitation due to linguistic predictability may be used more efficiently from individuals with stronger auditory-motor synchronization. A relevant question arising from this is: under what conditions is the impact of the motor system on speech comprehension the strongest? Previous work observed an impact of the motor system on speech comprehension in demanding listening conditions, such as listening to speech in noise (5,33). Our data suggests that this view might extend towards conditions of fast speech (which requires more experiments) or might interact with linguistic predictability.

Speech comprehension is a highly predictive process which is affected by different sources of predictions. Here we show that, while speech comprehension is optimal in a preferred auditory temporal regime, the motor-system provides a role for individual flexibility in continuous speech comprehension. Additionally, we report that the well-known faciliatory effects of linguistic predictability on speech comprehension interact with individual differences in the motor system. This sets the stage for future assessments of how predictions from these systems interact and under what circumstances the human brain relies more on one over the other.

## Ethics

Experiment 1 was approved by the ethics committee of the School of Social Sciences, University of Dundee, UK (No. UoD-SoSS-PSY-UG-2019-88). Procedures for Experiment 2 and the Control Experiment were approved by the Max Planck Society (No. 2017_12).

## Data accessibility

Preprocessed data and analysis scripts will be available via OSF upon publication.

## Acknowledgments

We thank Anna Broggi and Harry Watt for help with data acquisition (Exp. 1), Dr. Klaus Frieler for methodological advice, and Dr. Gregory Hickok for valuable comments on a previous version of the manuscript.

