## Supplementary Information for "Explaining flexible continuous speech comprehension from individual motor rhythms"

for

[2]Psychology
University of Dundee,
Dundee DD1 4HN, UK,

[3]Department of Psychology
University of Lübeck,
Lübeck, Germany,

[4]Center for Brain, Behavior, and Metabolism
University of Lübeck,
Lübeck, Germany,

[5]Department of Psychology
New York University,
New York, NY, USA

[6]Max Planck NYU Center for Language, Music, and Emotion
Frankfurt am Main, Germany, New York, NY, USA

[7]Ernst Strüngmann Institute for Neuroscience in Cooperation with Max Planck Society
Frankfurt am Main, Germany

Corresponding author:
Christina Lubinus
Department of Cognitive Neuropsychology
Max-Planck-Institute for Empirical Aesthetics
Grüneburgweg 14
D - 60322 Frankfurt am Main, Germany


#### Supplementary Notes

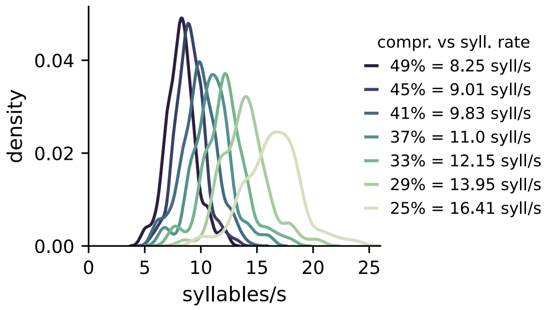


**Supplementary Fig. 1. Experiment 1 – Correspondence between syllabic rate and compression rate of speech stimuli.** Sentence stimuli were synthesized and then compressed to a percentage (49%, 45%, 41%, 37%, 33%, 29%) of their original duration, yielding sentences of different speech rates. Thus, the metric controlled for between conditions was compression rate. Since we were interested in the effect of syllabic rate on speech comprehension (not compression) and for better comparison with experiment 2, we computed the mean syllabic rate within each compression bin. To this end, we divided the number of syllables by the duration of the compressed stimulus which resulted in a distribution of syllabic rates. The distribution's means were defined as the corresponding syllabic rates. Importantly, the mapping between compression rate and syllabic rate is not unambiguous, as illustrated by the density plots. Color coding reflects the compression rates.

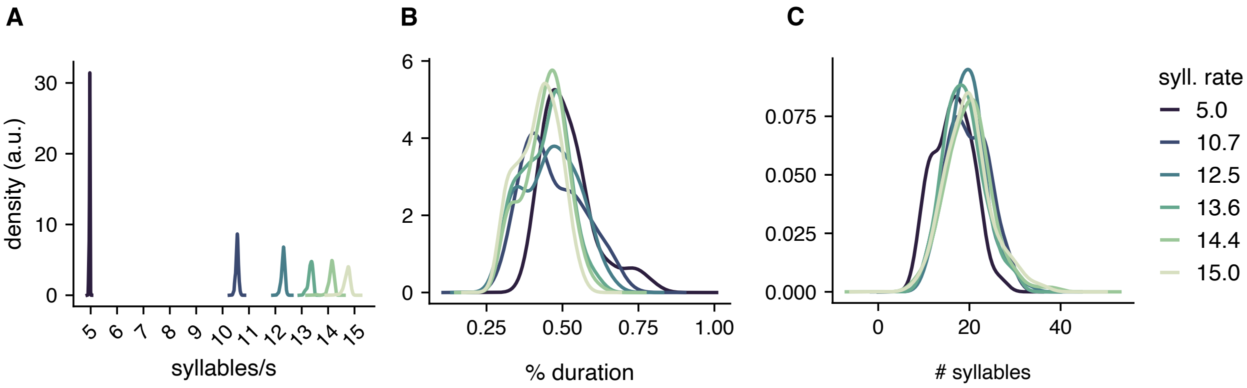


**Supplementary Fig. 2. Experiment 2 – distributional information for stimuli of speech comprehension task. A.** Syllabic rates. Speech stimuli were manipulated with respect to syllabic rate (5.0, 10.7, 12.5, 13.6, 14.4, 15.0 syllables/s), as visualized by separate distribution for each rate condition. Within rate conditions stimuli showed narrow distributions around the target rate. **B.** Compression rate. To assure comprehension performance was not confounded by systematic differences between compression strength between the syllabic rate conditions, the distribution of compression rates was kept as similar as possible across conditions (see Supplementary Methods 2). The density plot visualizes that the compression rates overlapped largely. **C.** Length of sentences. Across syllabic rates, the length of sentences (i.e. number of syllables) was similar as shown by overlapping distributions, suggesting that the rate conditions should not differ in effects related to sentence length (e.g., working memory load or complexity).

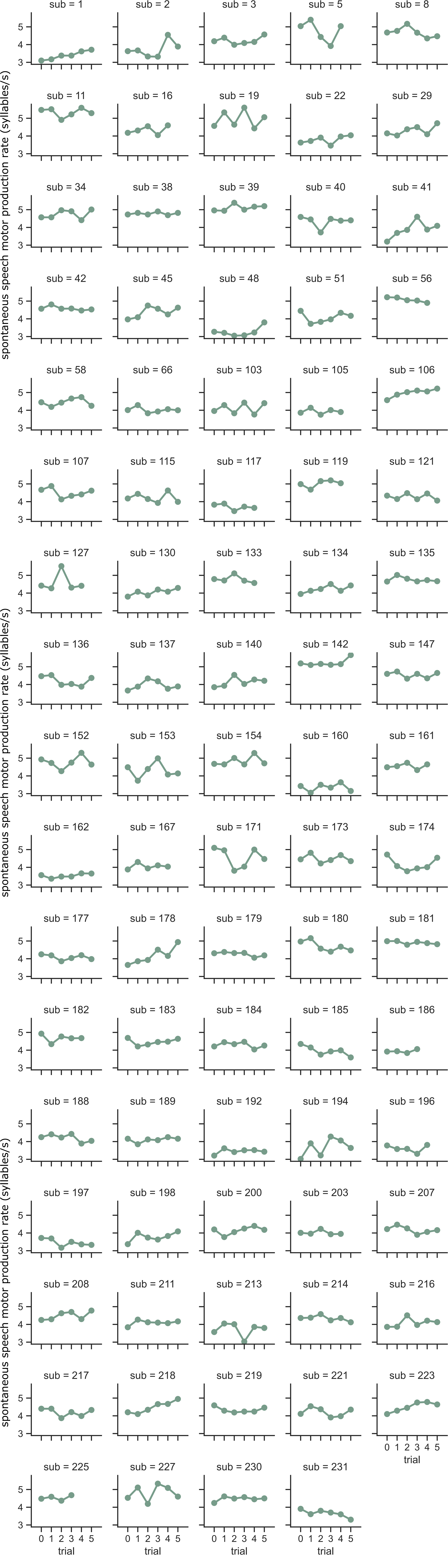


**Supplementary Fig. 3. Experiment 2 - Variation in spontaneous speech motor production rate.**
Single-trial estimates of spontaneous speech motor production rate, illustrated separately for each participant. The x-axis reflects the trial number; the y-axis reflects the spontaneous speech motor production rate (in syllables/s), as quantified across the duration of each trial (30 s). The final spontaneous speech motor production rate, used in the mixed model analysis, is the average across trials.

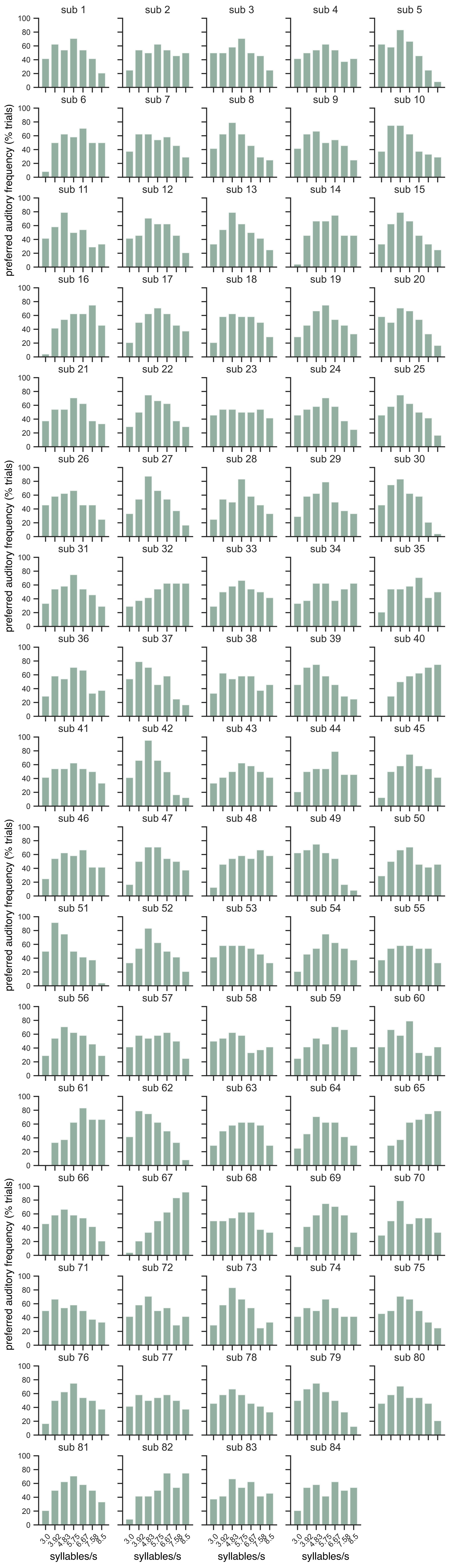


**Supplementary Fig. 4. Experiment 2 - Preferred auditory rates.**

Raw data from preferred auditory task, illustrated separately for each participant. The x-axis reflects the syllabic rate at which stimuli were presented; the y-axis reflects the percentage of all trials in which given syllabic rate was preferred as compared to the frequency it was contrasted with. This figure shows that most participants did show a preference for a syllabic rate, as indicated by a peaky distribution. However, some participants did not appear to have a strong preference for any syllabic rate as seen by means of a flat distribution.

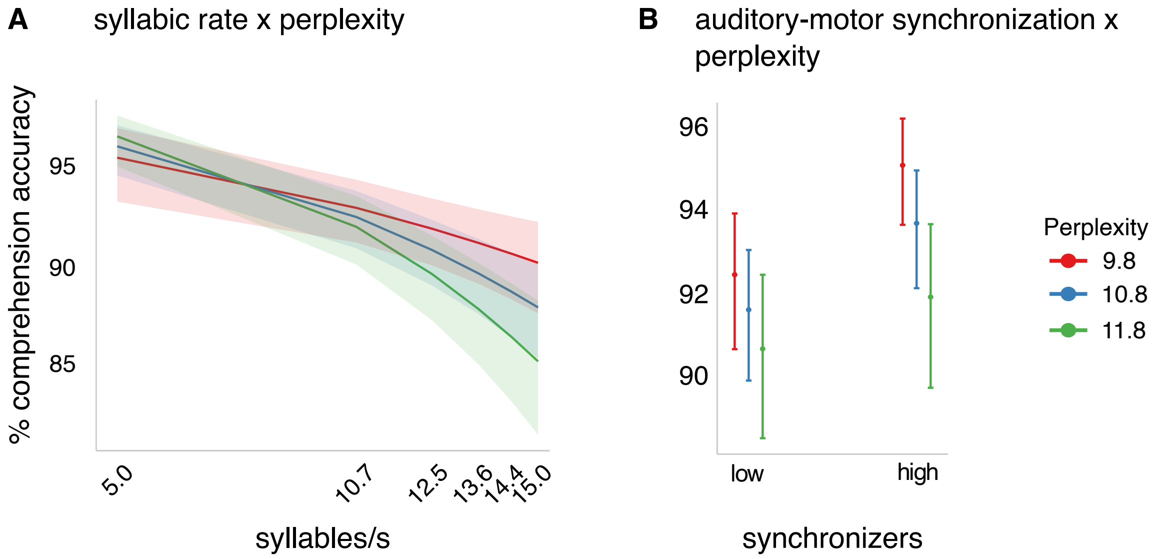


**Supplementary Figure 5. Experiment 2 – Interaction effects of *syllabic rate*, *auditory-motor synchronization*, and *perplexity*. A.** The generalized additive mixed model revealed a significant *syllabic rate* x *perplexity* interaction such that comprehension was best for sentences of highest predictability particularly at high demanding rates. **B.** We observed an interaction effect of *auditory-motor synchronization* x *perplexity,* suggesting a stronger comprehension gain as a function of predictability for high synchronizers, particularly when linguistic predictability is high.

**Supplementary Table 1. Experiment 2 – Sentence materials: Titles and authors of (audio)books.** List of sources from which stimuli for the tasks (speech comprehension and preferred auditory rate tasks) were constructed.

| Type | Author | Title | Source |
|---|---|---|---|
| Audiobook | James Weldon Johnson | The Autobiography Of An Ex-Colored Man | Lit2Go |
| Audiobook | Edith Wharton | Ethan Frome | Lit2Go |
| Audiobook | Frances Hodgson Burnett | A little princess | Librivox |
| Audiobook | Horace Walpole | Castle of Otranto | Lit2Go |
| Audiobook | Henry Ossian Flipper | The Colored Cadet At West Point | Lit2Go |
| PDF | Frances Hodgson Burnett | The Secret Garden | Lit2Go |
| PDF | Lucy Maud Montgomery | Anne of Green Gables | Lit2Go |
| PDF | Sinclair Lewis | The Job | Librivox |
| PDF | Booker T. Washington | Up from Slavery | Lit2Go |

**Supplementary Table 2. Experiment 2 – Speech production task.** Thematic questions used to facilitate natural speech production. Each item is representative of a different thematic category, as introduced by Alexandrou et al.[1].

| Category | Sentence/statement |
|---|---|
| Own life | What kind of hobbies do you have or have had during your life? |
| Preferences | What kinds of vacation trips do you like? |
| People | Describe a known artist, writer, or film director. Why do you find her/him interesting? |
| Culture/traditions | Describe a traditional Christmas holiday. |
| Society/politics | What do you know about garbage and recycling policies in your home country? |
| General knowledge | What do you know about skiing and snowboarding? |

**Supplementary Table 3. Experiment 1 – Predicting single-trial comprehension performance**

**Speech comprehension accuracy**

**Parametric coefficients**

| *Predictors* | *Estimate* | *std. Error* | *t-value* | *Pr(>ltl)* |
|---|---|---|---|---|
| Intercept | 54.720 | 2.188 | 25.01 | <0.001 |

**Approximate significance of smooth terms**

| | *edf* | *F-value* | *p-value* |
|---|---|---|---|
| Compression rate | 4.599 | 1241.436 | <0.001 |
| Speech motor prod. rate | 1.000 | 4.336 | 0.037 |
| sub | 30.231 | 17.077 | <0.001 |
| Trial-ID | 60.939 | 7.591 | <0.001 |

R-sq. (adj) = 76.1 Deviance explained = 77%
Observations = 2373

**Supplementary Table 4. Experiment 2 – Predicting single-trial comprehension performance.**

| | **comprehension_accuracy** | | | | |
|---|---|---|---|---|---|
| *Predictors* | *Odds Ratios* | *std. Error* | *CI (95%)* | *z-values* | *p* |
| (Intercept) | 11.27 | 1.18 | 9.18 – 13.84 | 23.14 | **<0.001** |
| syllabic rate | 0.65 | 0.04 | 0.57 – 0.73 | -6.70 | **<0.001** |
| synchronization (HIGH vs LOW) | 1.34 | 0.20 | 1.00 – 1.79 | 1.98 | **0.048** |
| speech motor prod. rate | 1.19 | 0.09 | 1.04 – 1.37 | 2.45 | **0.014** |
| pref. auditory rate | 1.14 | 0.08 | 0.99 – 1.31 | 1.80 | 0.072 |
| working memory | 1.20 | 0.09 | 1.04 – 1.39 | 2.52 | **0.012** |
| perplexity | 0.84 | 0.04 | 0.76 – 0.94 | -3.19 | **0.001** |
| probability target1 | 0.93 | 0.03 | 0.88 – 0.99 | -2.23 | **0.026** |
| probability target2 | 0.92 | 0.03 | 0.85 – 0.99 | -2.31 | **0.021** |
| compression | 1.21 | 0.06 | 1.10 – 1.33 | 3.87 | **<0.001** |
| sentence length | 0.61 | 0.03 | 0.55 – 0.68 | -9.24 | **<0.001** |
| target distance | 1.48 | 0.05 | 1.37 – 1.59 | 10.60 | **<0.001** |
| syllabic rate * synchronization | 0.97 | 0.07 | 0.84 – 1.10 | -0.52 | 0.602 |
| **Random Effects** | | | | | |
| $\sigma^2$ | 3.29 | | | | |
| $\tau_{00\ file}$ | 0.74 | | | | |
| $\tau_{00\ sub}$ | 0.36 | | | | |
| $\tau_{11\ sub.scale(freq)}$ | 0.02 | | | | |
| $\rho_{01\ sub}$ | 0.00 | | | | |
| ICC | 0.25 | | | | |
| $N_{sub}$ | 82 | | | | |
| $N_{file}$ | 495 | | | | |
| Observations | 19680 | | | | |
| Marginal $R^2$ / Conditional $R^2$ | 0.139 / 0.357 | | | | |

**Supplementary Table 5. Experiment 2 – Predicting single-trial comprehension performance (with FDR-correction)**

| | comprehension_accuracy | | | | |
|---|---|---|---|---|---|
| *Predictors* | *Odds Ratios* | *std. Error* | *CI (95%)* | *z-values* | *p* |
| (Intercept) | 11.27 | 1.18 | 9.18 – 13.84 | 23.14 | **<0.001** |
| syllabic rate | 0.65 | 0.04 | 0.57 – 0.73 | -6.70 | **<0.001** |
| synchronization (HIGH vs LOW) | 1.34 | 0.20 | 1.00 – 1.79 | 1.98 | 0.057 |
| speech motor prod. rate | 1.19 | 0.09 | 1.04 – 1.37 | 2.45 | **0.023** |
| pref. auditory rate | 1.14 | 0.08 | 0.99 – 1.31 | 1.80 | 0.078 |
| working memory | 1.20 | 0.09 | 1.04 – 1.39 | 2.52 | **0.022** |
| perplexity | 0.84 | 0.04 | 0.76 – 0.94 | -3.19 | **0.003** |
| probability target1 | 0.93 | 0.03 | 0.88 – 0.99 | -2.23 | **0.034** |
| probability target2 | 0.92 | 0.03 | 0.85 – 0.99 | -2.31 | **0.030** |
| compression | 1.21 | 0.06 | 1.10 – 1.33 | 3.87 | **<0.001** |
| sentence length | 0.61 | 0.03 | 0.55 – 0.68 | -9.24 | **<0.001** |
| target distance | 1.48 | 0.05 | 1.37 – 1.59 | 10.60 | **<0.001** |
| syllabic rate * synchronization | 0.97 | 0.07 | 0.84 – 1.10 | -0.52 | 0.602 |
| **Random Effects** | | | | | |
| $\sigma^2$ | 3.29 | | | | |
| $\tau_{00\ file}$ | 0.74 | | | | |
| $\tau_{00\ sub}$ | 0.36 | | | | |
| $\tau_{11\ sub.scale(freq)}$ | 0.02 | | | | |
| $\rho_{01\ sub}$ | 0.00 | | | | |
| ICC | 0.25 | | | | |
| $N_{sub}$ | 82 | | | | |
| $N_{file}$ | 495 | | | | |
| Observations | 19680 | | | | |
| Marginal $R^2$ / Conditional $R^2$ | 0.139 / 0.357 | | | | |

**Supplementary Table 6. Experiment 2 – Predicting single-trial comprehension performance including a 3-way interaction term of syllabic rate x synchronization x perplexity**. Output of generalized mixed-effects model, showing predictions of single-trial speech comprehension when controlling for word-order effects of the target words.

| | comprehension_accuracy | | | | |
|---|---|---|---|---|---|
| *Predictors* | *Odds Ratios* | *std. Error* | *CI (95%)* | *z-values* | *p* |
| (Intercept) | 11.12 | 1.16 | 9.06 – 13.65 | 23.01 | **<0.001** |
| syllabic rate | 0.66 | 0.04 | 0.58 – 0.75 | -6.37 | **<0.001** |
| synchronization (HIGH vs LOW) | 1.35 | 0.20 | 1.01 – 1.81 | 2.04 | **0.041** |
| perplexity | 0.89 | 0.05 | 0.80 – 1.00 | -1.99 | **0.046** |
| speech motor prod. rate | 1.19 | 0.09 | 1.04 – 1.37 | 2.45 | **0.014** |
| pref. auditory rate | 1.14 | 0.08 | 0.99 – 1.31 | 1.80 | 0.072 |
| working memory | 1.20 | 0.09 | 1.04 – 1.39 | 2.52 | **0.012** |
| probability target1 | 0.93 | 0.03 | 0.88 – 0.99 | -2.20 | **0.028** |
| probability target2 | 0.91 | 0.03 | 0.85 – 0.98 | -2.41 | **0.016** |
| compression | 1.22 | 0.06 | 1.10 – 1.34 | 4.00 | **<0.001** |
| sentence length | 0.61 | 0.03 | 0.55 – 0.68 | -9.17 | **<0.001** |
| target distance | 1.47 | 0.05 | 1.37 – 1.58 | 10.58 | **<0.001** |
| syllabic rate * synchronization | 0.94 | 0.07 | 0.82 – 1.08 | -0.86 | 0.392 |
| syllabic rate * perplexity | 0.88 | 0.05 | 0.80 – 0.98 | -2.43 | **0.015** |
| synchronization * perplexity | 0.86 | 0.04 | 0.78 – 0.95 | -2.96 | **0.003** |
| syllabic rate * synchronization * perplexity | 1.09 | 0.06 | 0.98 – 1.21 | 1.62 | 0.106 |
| **Random Effects** | | | | | |
| $\sigma^2$ | 3.29 | | | | |
| $\tau_{00\ file}$ | 0.73 | | | | |
| $\tau_{00\ sub}$ | 0.56 | | | | |
| $\tau_{11\ sub.freq}$ | 0.00 | | | | |
| $\rho_{01\ sub}$ | -0.60 | | | | |
| ICC | 0.28 | | | | |
| $N_{sub}$ | 82 | | | | |
| $N_{file}$ | 495 | | | | |
| Observations | 19680 | | | | |
| Marginal $R^2$ / Conditional $R^2$ | 0.139 / 0.382 | | | | |

**Supplementary Table 7. Control experiment – Predicting single-trial comprehension performance including the word order index**. Output of generalized mixed-effects model, showing predictions of single-trial speech comprehension when controlling for word-order effects of the target words.

| | comprehension_accuracy | | | | |
|---|---|---|---|---|---|
| *Predictors* | *Odds Ratios* | *std. Error* | *CI (95%)* | *z-values* | *p* |
| (Intercept) | 11.25 | 1.18 | 9.17 – 13.82 | 23.13 | **<0.001** |
| syllabic rate | 0.65 | 0.04 | 0.57 – 0.73 | -6.69 | **<0.001** |
| synchronization (HIGH vs LOW) | 1.34 | 0.20 | 1.01 – 1.80 | 2.01 | **0.045** |
| speech motor prod. rate | 1.19 | 0.09 | 1.03 – 1.37 | 2.42 | **0.016** |
| pref. auditory rate | 1.14 | 0.08 | 0.99 – 1.31 | 1.80 | 0.072 |
| working memory | 1.20 | 0.09 | 1.04 – 1.39 | 2.47 | **0.013** |
| perplexity | 0.84 | 0.04 | 0.76 – 0.94 | -3.19 | **0.001** |
| probability target1 | 0.93 | 0.03 | 0.88 – 0.99 | -2.19 | **0.028** |
| probability target2 | 0.92 | 0.03 | 0.85 – 0.99 | -2.30 | **0.022** |
| compression | 1.21 | 0.06 | 1.10 – 1.33 | 3.89 | **<0.001** |
| sentence length | 0.61 | 0.03 | 0.55 – 0.68 | -9.23 | **<0.001** |
| target distance | 1.48 | 0.05 | 1.37 – 1.59 | 10.62 | **<0.001** |
| word order | 0.96 | 0.03 | 0.90 – 1.02 | -1.23 | 0.219 |
| syllabic rate * synchronization | 0.97 | 0.07 | 0.84 – 1.10 | -0.52 | 0.606 |
| **Random Effects** | | | | | |
| $\sigma^2$ | 3.29 | | | | |
| $\tau_{00\ file}$ | 0.74 | | | | |
| $\tau_{00\ sub}$ | 0.55 | | | | |
| $\tau_{11\ sub.freq}$ | 0.00 | | | | |
| $\rho_{01\ sub}$ | -0.59 | | | | |
| ICC | 0.28 | | | | |
| $N_{sub}$ | 82 | | | | |
| $N_{file}$ | 495 | | | | |
| Observations | 19680 | | | | |
| Marginal $R^2$ / Conditional $R^2$ | 0.134 / 0.379 | | | | |

### Supplementary Methods

#### Participants

Experiment 1

Participation was voluntary and participants had the chance of winning a £25 voucher by participating in a prize draw. We recruited participants through opportunity sampling at the University of Dundee. All participants were native English speakers (N = 34, female = 18, male = 14, non-binary = 2, age: M = 22.12, SD = 1.87), right-handed and reported normal hearing, as well as no neurological or psychological disorders. The experiment complied with the Declaration of Helsinki and was approved by the ethics committee of the School of Social Sciences, University of Dundee, UK (No. UoD-SoSS-PSY-UG-2019-88).

The experimental tasks were presented using Psychtoolbox (version 3.0.16) for MATLAB (version R2017b) on a Windows computer. Participants were equipped with non-wireless DRACO HS-880 headphones (Creative) with an integrated microphone to record all speech signals.

Experiment 2

Participants were native speakers of North-American English, born in the United States or Canada, recruited from the online portal “Prolific” (https://www.prolific.com/). The following criteria were used for Prolific’s participants prescreening: normal or corrected-to-normal vision, no hearing issues, no prior or current psychological or neurological diseases, aged 18-45 years. Furthermore, using Prolific’s compliance metrics, we only included participants with a minimum approval rate of 90% in previous experiments and a minimum of 50 previous submissions. The final sample included 82 participants (37 Females; age: *M* = 28.6 years; *SD* = 6.3), 36 High and 46 Low synchronizers.

Experimental tasks were presented in the web browser using JsPsych (6.1.0) and the experiment was hosted on an in-house Jatos (3.5.5) server. The experiment complied with the Declaration of Helsinki and the procedures were approved by the Ethics Council of the Max Planck Society (2017_12).

Control experiment

A new set of participants (N = 39, 13 per stimulus list, female = 10, age: M = 29.1, STD = 10.1) was recruited from Prolific (same inclusion criteria as in Experiment 2, enriched only by the criterion of not having participated in the previous study). Experimental setup and ethics approval are identical to Experiment 2.

#### Stimulus selection, recording, and processing

Experiment 1 – Speech comprehension task

During stimulus generation, first, sentences (between 5 to 8 words) were generated using the online tool SKELL (https://skell.sketchengine.co.uk/run.cgi/skell). Second, the sentences were synthesized using Google Cloud’s text2speech (male voice ’en-GB-Wavenet-B’, https://cloud.google.com/text-to-speech) to generate audio files at a sampling rate of 44,100 Hz and 2dB volume gain. The advantage of this text2speech algorithm is its ability to produce human-like speech generating stimuli consistent in speech rate and loudness. Finally, the synthesized audio files were digitally compressed using the Pitch Synchronous Overlap and Add (PSOLA) algorithm[2] implemented in Praat (6.0.18)[3].

Importantly, speech rate was manipulated by means of compression rate (percentage of stimulus duration), that is, compression rate varied between different syllabic rates. For stimulus definition/generation and the statistical analysis, stimuli were grouped based on compression rate. For visualization and easier comparison with the second experiment, we transformed compression rate into *syllabic rate* (syllables/s). The mapping between compression and syllabic rate is not unambiguous in that any given compression rate contained distinct syllabic rates (for correspondence between both measures see Supplementary Fig. 1).

Experiment 1 – Reading excerpt for speech production task

Dennis was different. When he looked in the mirror, he saw an ordinary twelve-year-old boy. But he felt different – his thoughts were full of colour and poetry, though his life could be very boring. The story I am going to tell you begins here, in Dennis's ordinary house on an ordinary street in an ordinary town. His house was nearly exactly the same as all the others in the street. One house had double glazing, another did not. One had a gravel drive, another had crazy paving. One had a Vauxhall Cavalier in the drive, another a Vauxhall Astra. Tiny differences that only really pointed out the sameness of everything. It was all so ordinary, something extraordinary just had to happen. Dennis lived with his dad – who did have a name, but Dennis just called him Dad, so I will too – and his older brother John, who was fourteen. Dennis found it frustrating that his brother would always be two years older than him, and bigger, and stronger. Dennis's mum had left home a couple of years ago. Before that, Dennis used to creep out of his room and sit at the top of the stairs and listen to his mum and dad shout at each other until one day the shouting stopped. She was gone.

Experiment 2 – Sentence materials for speech comprehension and preferred auditory tasks

When studying the comprehension of accelerated speech, the syllabic rate of stimuli is usually manipulated by digital compression. A consisting finding is that comprehension or intelligibility deteriorates as the syllabic rate of sentences increases[4–7]. In these studies, the degree of compression and the syllabic rate of sentences are typically correlated, such that faster sentences are compressed more heavily. Importantly, digital acceleration not only alters speech rate but also the acoustic properties of a signal[8]. Considering the relevance of acoustic features for speech intelligibility (e.g. edges[9]), in Experiment 2 we carefully controlled for potential compression artifacts by balancing compression factors across syllabic rates. To this end, we constructed a set of stimuli with a broad range of original speech rates. As speech rate in audiobooks, and natural speech more generally, is robustly centered around 4.5 Hz[10,11], part of or stimuli were recorded by speakers speaking as fast as possible while maintaining proper articulation.

All sentence materials were sourced from books and audiobooks that are freely available on LibriVox (https://librivox.org/) or Lit2Go (https://etc.usf.edu/lit2go/) (see Supplementary Table 1). The final stimulus set was drawn from five audiobooks and four books for which the recordings were performed by native speakers of North-American English at the Max Planck Institute for Empirical Aesthetics.

Recordings were performed in a sound-attenuated recording booth using MatLab R2017a (Version: 9.2.0.538062) on a Windows 7 Pro (64-bit) and a Neumann U87i studio microphone (https://en-de.neumann.com/u-87-ai) and digitized to a sampling rate of 44 kHz. Speakers spoke them at two speeds: at a normal, natural pace and as fast as possible (while maintaining proper articulation). A total of 788 individual sentences was acquired. In addition to the fast recordings, one batch of stimuli was recorded from a speaker who was instructed to speak as slowly as possible. Processing of all sound files (audio books and recordings) was performed using Praat (6.0.40)[3]. Long pauses (> 300ms) were removed to avoid wrong estimates for syllables per second. After compression and expansion of the audio files, all final stimuli were matched for root mean square (RMS) amplitude (69 dB).

Experiment 2 – Stimulus lists for speech comprehension and preferred auditory tasks

From the recordings (N = 10 speakers), we created three stimulus lists for the speech comprehension (N = 240 sentences) and the auditory rate tasks (N = 49 individual sentences) by randomly drawing sentences (without replacement) from the total pool of 788 sentences. The range of original syllabic rates was 1.97-9.71 syllables/s. Within each set of stimulus lists (i.e. speech comprehension and auditory rate tasks) no sentences were repeated. Original speech recordings were time-compressed and time-expanded using the Pitch Synchronous Overlap and Add (PSOLA)[2] algorithm.

The original sentences were compressed/expanded to the following syllabic rates: speech comprehension task: 5.00, 10.69, 12.48, 13.58, 14.38, 15.00; preferred auditory rate task: 3.00, 3.92, 4.83, 5.75, 6.67, 7.58, 8.50. Syllabic rate conditions were matched for sentence length (number of syllables), compression rate, position of target words, number of speakers (Supplementary Fig. 2). Maximal compression and expansion were constrained (compression: factor of 3; expansion: factor of 2). For the slowest condition (5 syllables/s) the variation in speakers is lower because only one speaker could be recorded (and no further recordings were possible due to lockdown).

Experiment 2 – Stimuli for digit span test

The digits 0-9 were synthesized using the Mac OSX text-to-speech application (voice Anna). Using Audacity, each digit was concatenated with silence such that digit and silence amounted to a duration of one second. Digit spans[12] were created by concatenating the corresponding digits using Audacity so that the digits occurred at a rate of one digit per second.

#### Experimental tasks

Speech comprehension task

Experiment 1

Participants performed an intelligibility task. On each trial (N = 70), a sentence was presented through headphones and participants verbally repeated all perceived words. Responses were recorded. Sentences were presented at various syllabic rates (8.2, 9.0, 9.8, 11.0, 12.1, 14.0, 16.4.) and grouped into blocks according to syllabic rates. While the syllabic rates were presented in the same (descending) manner to all participants, sentences were randomly assigned to each block/syllabic rate.

Experiment 2

Participants performed a word-order task in which they listened to one sentence per trial (N = 240), followed by the presentation of two words from the sentence on screen. Participants indicated via button press which word they heard first.

The sentences were presented at various syllabic rates: 5.00, 10.69, 12.48, 13.58, 14.38, 15.00. For each syllabic rate 40 sentences were presented, amounting to a total of 240 sentences. Trials were grouped into blocks of 20 sentences, allowing for self-paced breaks between blocks. We randomized sentence order and syllabic rates across participants. The maximal response time was 5000 ms. The inter-trial interval was jittered (uniform distribution between 1000 and 1500 ms). Prior to task begin, participants were familiarized with the task by 1) listening to one stimulus played at all possible frequencies (without task) and 2) performing three practice trials.

Auditory rate task (only Experiment 2)

Participants performed a 2IFC task, in which a reference and a comparison stimulus were presented. Within a trial, reference and comparison stimulus were constructed from the same sentence – they only differed with regard to syllabic rate (3.00, 3.92, 4.83, 5.75, 6.67, 7.58, 8.50). Reference and comparison stimulus were separated by an inter-stimulus interval (uniform distribution between 500 and 1000 ms) and trials by a randomized inter-trial interval (uniform distribution between 1000 and 1500 ms).

To ensure participants were engaged in the task, we added catch trials (one trial for each reference frequency, i.e., 7 trials). Catch stimuli were manipulated such that one syllable was repeated three times in both the reference and comparison stimuli. Participants were instructed to respond to catch trials by pressing "t", instead of "x" or "m", and were informed that poor performance would lead to an exclusion from the experiment. Prior to the main task, participants were familiarized with the task by performing three practice trials for the main task, as well as one practice trial for the catch trials.

Digit span test (only Experiment 2)

The digits 0-9 were synthesized using the Mac OSX text-to-speech application (voice Anna). Using Audacity, each digit was concatenated with silence such that digit and silence amounted to a duration of one second. Digit spans[12] were created by concatenating the corresponding digits using Audacity so that the digits occurred at a rate of one digit per second.

The test comprised seven levels (3 to 9 digits) with two items at each level. The procedure started with the shortest digit spans (3 digits) and stopped as soon as participants failed to correctly repeat both spans belonging to one length or when the two longest digit spans (9 digits) were finished.

Procedure

Experiment 1 was conducted in the laboratory at Dundee University. Participants were seated in front of a computer in a quiet experimental room. Prior to experiment start, participants completed a demographic questionnaire. Upon giving informed consent, the speech comprehension task was completed first, followed by the speech production task.

Experiment 2 entailed three sessions all of which were conducted online. Overall, the protocol contained 5 tasks (3 main tasks: comprehension task, auditory rate task, speech production task; 2 further tasks: SSS-test, digit span test) and a questionnaire. In each session, one of the three main tasks was completed. The order was randomized across participants. To assure participants wore headphones, a headphone screening test was performed in the be-ginning of each session (55). At the end of the last session, the SSS test, digit span test, and the questionnaire were conducted (see Supplementary Fig. 3).

##### Exclusion criteria (Experiment 2)

Only complete datasets (all three sessions completed) were considered for analysis. From participants with complete datasets, participants were excluded if (a) comprehension performance was 2 *SD* below mean performance at the baseline rate (5 syll/s) in the speech comprehension task, (b) detection of catch trials was below 75% in the auditory rate task, (c) participants did not whisper but speak normally/loudly during the SSS test, (d) poor audio recordings.

##### Computing word predictability and perplexity.

To account for variation in sentence predictability in the comprehension task, we created a recurrent neural network (RNN) language model which assigns probabilities to sequences (for similar analysis see [13]). RNNs, especially when using long short-term memory (LSTM)[14], are well suited to approximate language processing because they can incorporate past input (e.g., words) into the prediction of a current word[15], just as done during natural language processing in humans. The model was created to solve the task of predicting each word in the sentence based on the previous word or words. For the training, sentences were taken from a variety of books, to then predict word probabilities for all sentences in the stimulus materials used in our experiment. For this analysis we used Keras[16] with a Tensorflow[17] backend.

The training dataset was curated from 101 freely available classic books and underwent a cleaning procedure (removing punctuation, lower casing all words). We aimed at creating training materials as similar as possible to our stimuli. Therefore, books were selected from the same authors as the stimulus materials where possible, as well as different authors but similar topics and genres. Sentences were selected as training material only if their length matched the stimulus materials (8-25 words). The final data set comprised 252.377 sentences, constructed from 38.195 unique words. Prior to training, the curated data set was split into training and evaluation sets (75 to 25%).

For the RNN analysis, we worked off the scripts provided by ten Oever & Martin[13]. Our RNN consisted of 3 layers: a pretrained embedding layer using Google's word2vec embeddings (https://code.google.com/archive/p/word2vec/), an LSTM layer with a tanh activation (300 hidden units) and a dense output layer (softmax activation). The model was trained using the following hyperparameters: batch size = 32, epochs = 100, Adam optimization with a learning rate of 0.001, and regularization to prevent overfitting (dropout of 0.2 for recurrent and output layers and L2 regularization of 0.001). Given the multi-class classification problem (which word out of 38.195 words is most likely to follow), we implemented a sparse categorical cross-entropy loss function.

Upon training, the trained model was used to evaluate and predict sentence predictability of all stimulus sentences (test set, N = 495) used in the comprehension task. From the RNN predictions of the test set, we derived two measures: target word probability (for both target words) and predictability of the whole sentence. To quantifying each sentences' predictability[18,19], we derived one value per sentence from the single-word probabilities. This so-called perplexity is the most common intrinsic evaluation metric of language models[15,20,21]. It is computed as the inverse of the mean probability of a sentence weighted by sentence length[18], i.e. lower perplexity values equal higher sentence predictability.

#### Control analyses

Correlation of digit span and preferred auditory rate/spontaneous speech motor production rate

Digit span and speech motor production rate: rho = 0.064, p = 0.569

Digit span and preferred auditory rate: rho = 0.040, p = 0.728

Correlation of preferred auditory rate and spontaneous speech motor production rate

rho = -0.052, p = .724
